## Supplemental material for "EEG-fMRI in awake rat and whole-brain simulations show decreased brain responsiveness to sensory stimulations during absence seizures"

**Figure supplements**

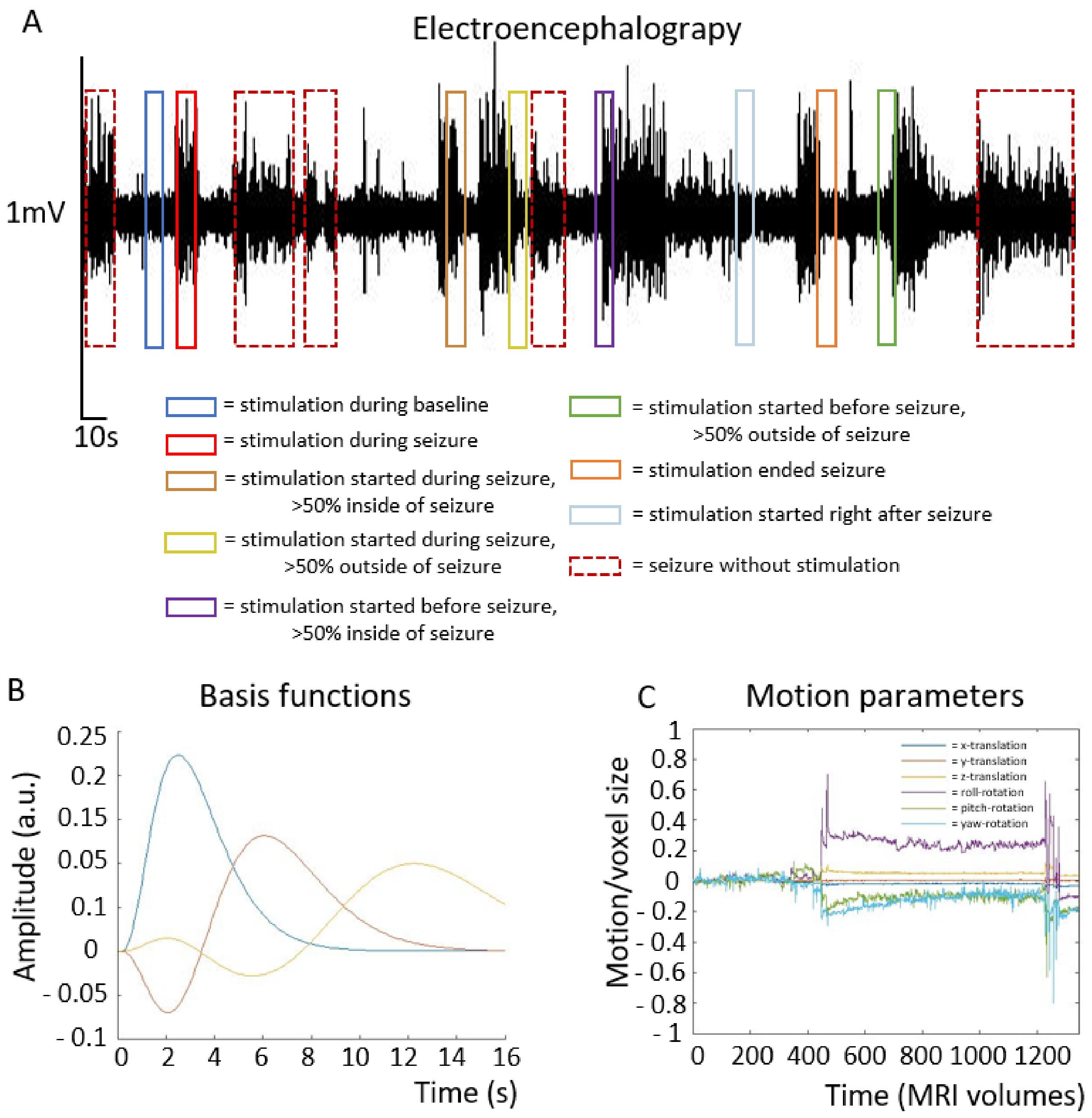


**Figure 1–figure supplement 1. Example of EEG trace and illustration of stimulation and seizure block paradigms used as SPM inputs.** Inputs (A) were convolved with 3$^{rd}$ order gamma functions (B), which was selected as basis functions, accounting for temporal and dispersion differences in hemodynamic responses. Example of translational and rotational motion parameters from one animal (C). Motion parameters were used as nuisance regressors and not convolved with a basis function.

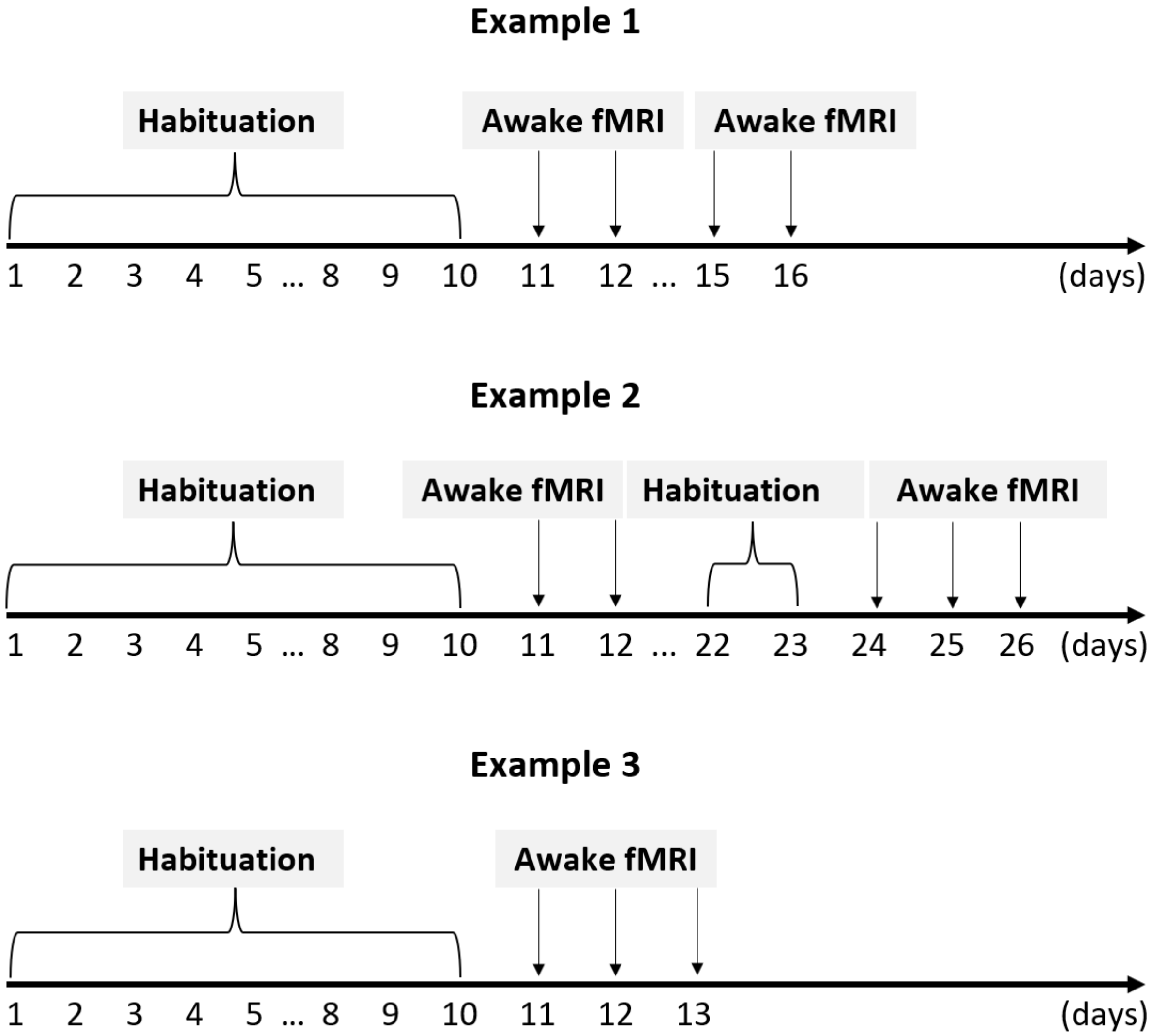


**Figure 1–figure supplement 2. Habituation and imaging schedules for three illustrative rats.** After 8 days of habituation**,** 3-5 fMRI experiment were conducted within a 1–3-week period for each rat. In case rats were re-imaged more than 1 week after the preceding experiment, an additional 2-day habituation period was conducted.

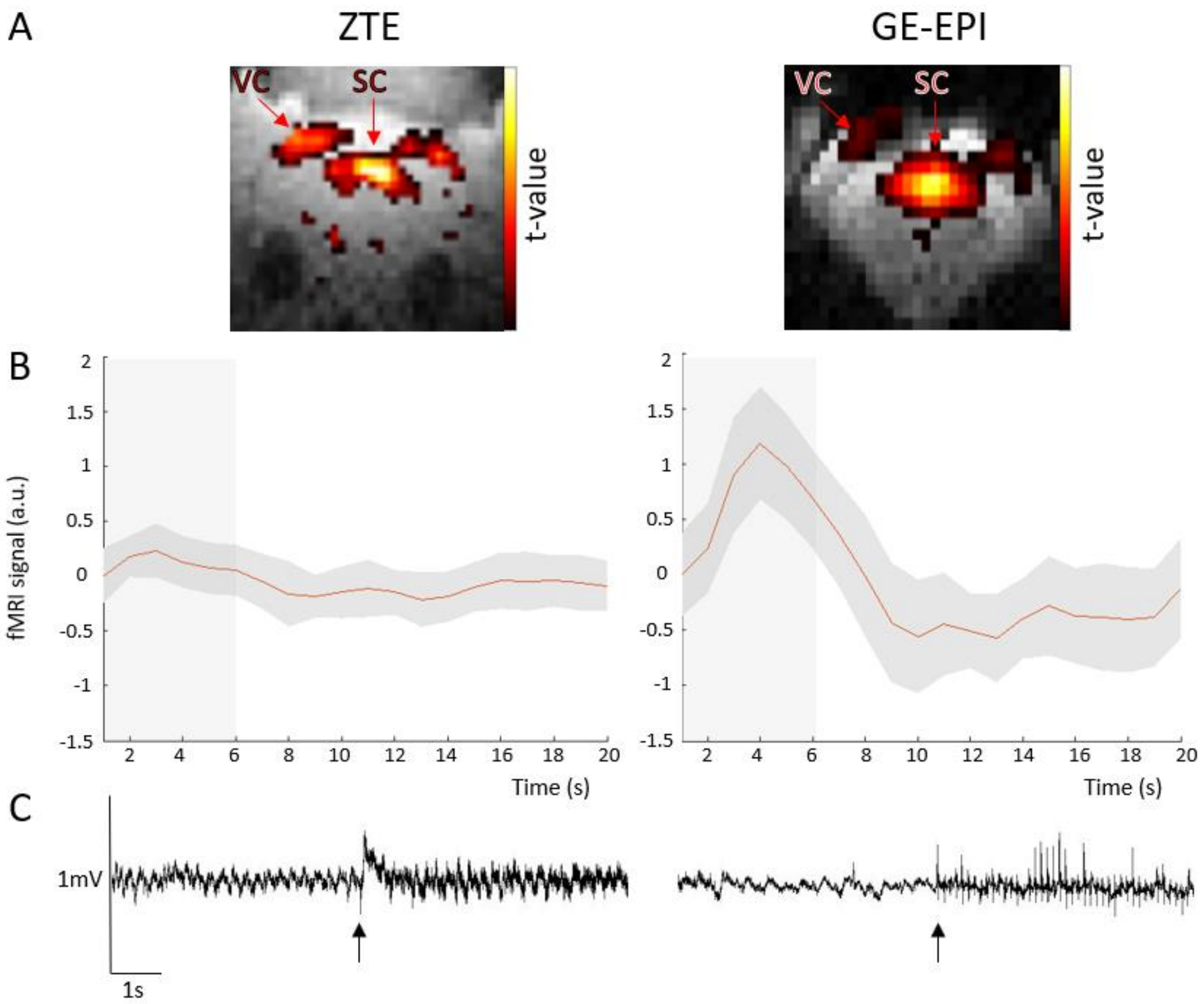


**Figure 2–figure supplement 1. ZTE and gradient-echo-EPI (GE-EPI) data collected during visual stimulus experiment, and EEG trace illustrating MRI gradient artefacts, from an example rat in preliminary study.** Illustration of image quality, and statistical t-contrast maps to visual stimulation (A), functional contrast (B) and effect of EEG artefact (C) for both imaging sequences. Although ZTE exhibits lower functional contrast compared to EPI-sequence, it offered a better spatial brain coverage with less image distortions, thus yielding far better stimulation induced maps. ZTE also caused relatively less artificial noise on EEG signal, keeping both amplitude of the signal and frequencies relatively more intact, which improved live detection of absence seizures. Grey box illustrates stimulation period. Arrows on EEG signal mark the beginning of each fMRI sequence.

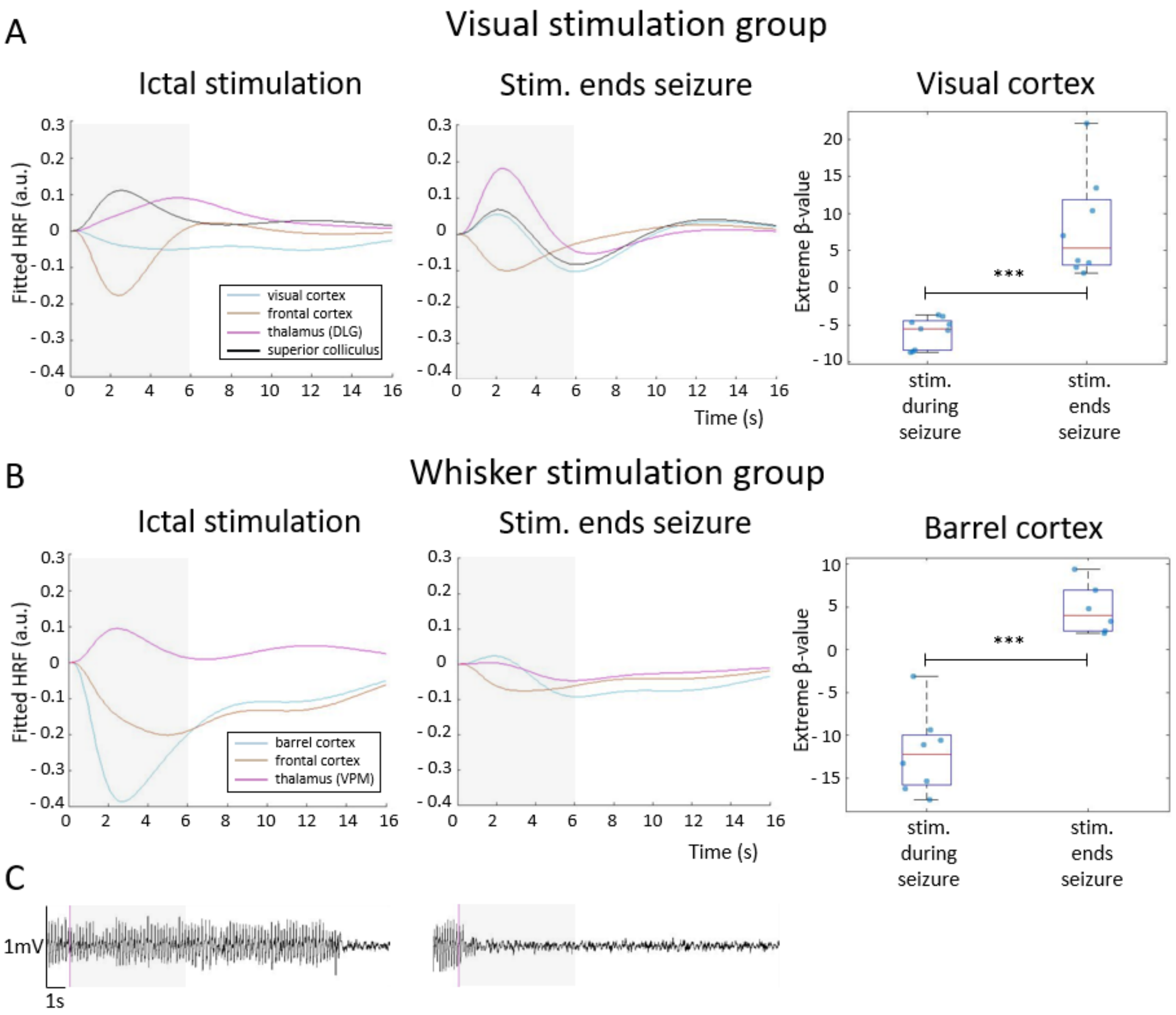


**Figure 4–figure supplement 1. Hemodynamic response functions to stimulation during an ictal period and during a condition when stimulation ended a seizure, for visual stimulation (A) and whisker stimulation (B) groups.** HRF was calculated in selected ROI, belonging to visual or somatosensory area, by multiplying gamma basis functions (Figure 1–figure supplement 1B) with their corresponding average beta values over a ROI and taking a sum of these values. For statistical comparison, extreme beta-values over a ROI were calculated and values between two states were compared with a two-sample t-test. Scatter plots represent mean ± SD and each blue dot corresponds to the extreme beta-values observed during individual fMRI sessions. A representative period from EEG trace, with a stimulation onset marked (purple), in both conditions is also illustrated (C). *** = $p < 0.001$. Grey box illustrates stimulation period.

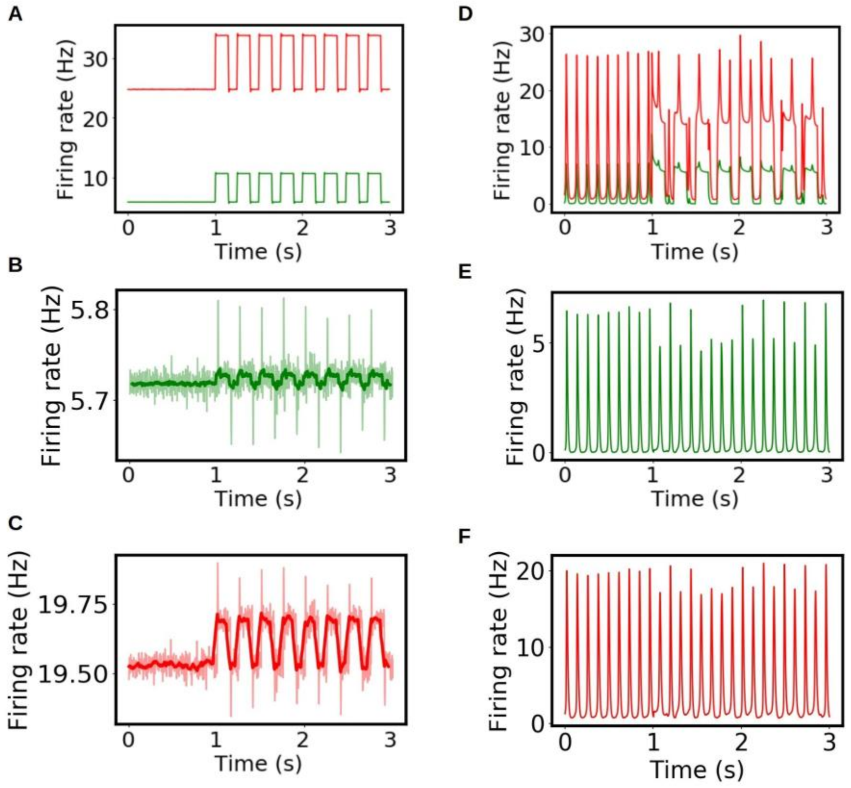


**Figure 6–figure supplement 1. Single region activity during stimulation in whole brain simulation.** We show the results of the simulations during inter-ictal (A, B, C) and ictal periods (D, E, F). The stimulus is applied in the primary visual cortex (A, D) and propagates to the connected regions. In panels B, C, E and F we show the excitatory (green) and inhibitory (red) activity in the mediolateral visual area during inter-ictal (B, C) and ictal (E, F) periods. For a better visualization the rolling average is shown in dark green and dark red respectively.

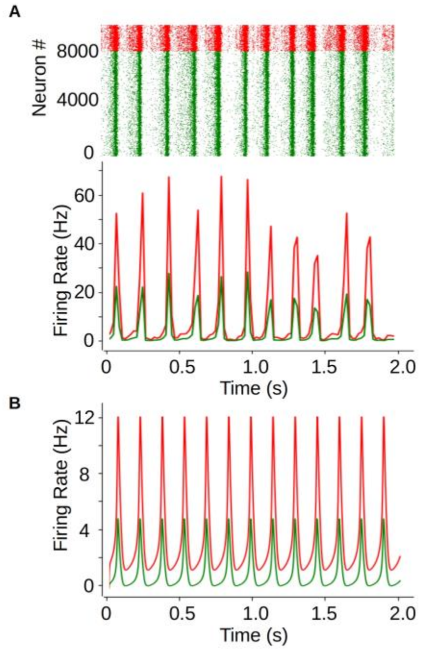

**Figure 6–figure supplement 2. Comparison of the SWD dynamics in the mean-field model and a spiking-neural network of AdEx neurons.** A) Raster plot (top) and mean firing rate (bottom) from an SWD type of dynamics obtained from the spiking- network simulations. The network is made of 8000 excitatory neurons and 2000 inhibitory neurons. Neurons in the network are randomly connected with a probability p=0.05 for inhibitory-inhibitory and excitatory-inhibitory connections, and p=0.06 for excitatory-excitatory connections. Cellular parameters correspond to the ones used in the mean-field, with spike-triggered adaptation for excitatory neurons set to b=200pA. We show the results for excitatory (green) and inhibitory (red) neurons. B) Mean-firing rate obtained from a single mean-field model. We see that, although the amplitude of oscillations is larger in the spiking-network, the mean-field can correctly capture the general dynamics and frequency of the oscillations.

### Appendix

#### Physiologic and methodologic considerations

As $P_{O2}$, $P_{CO2}$, and arterial blood pressure were not measured during the fMRI, there is a possibility that they affect the ZTE-fMRI readout, which is sensitive to blood flow changes. However, as animals were awake and given the fine cerebral autoregulation, blood flow values can be expected to be in the normal range. Another concern is whether blood flow remains stable during the seizure e.g. increases to a level that could hinder stimulus detection. However, previous doppler flowmetry studies have shown rather decreased than increased blood velocities during an absence seizure in both humans and rats (Bode, 1992; De Simone et al., 1998; Neblig et al., 1996). Also, our HRF results measured during a seizure in absence of stimulation suggest a similar finding of reduced cerebral blood flow (CBF) especially in cortical regions. Even then, we cannot rule out the possibility of CBF changes influencing the results, especially if there are CBF changes caused by non-neuronal origins. We note a caution that presented maps and time courses showing fMRI changes from visual or whisker stimulation during seizures may contain a mixture of both sensory stimulation-related signals and seizure-related signals. To minimize this contamination in the linear model used, we considered both stimulation and seizure-only states as regressors of interest and used seizure-only responses as nuisance regressors to account for error variance. Thereby, the effects caused by the stimulation should be separated as much as possible from the effects caused by the seizure itself.

As the used awake habituation and imaging protocol didn’t allow us to avoid the usage of isoflurane during the preparation steps, we cannot rule out the possible effect of using repetitive anesthesia on brain function. However, duration (~15 min) and concentration of anesthesia (~1.5%) during these steps were still moderate, whereas extended durations (1-3 h) of either single or repetitive isoflurane exposures have been used in previous studies where long-term effects on brain function have been observed (Long et al., 2016; Stenroos et al., 2021). Moreover, there was a 5-15 min waiting period between the cessation of anesthesia and initiation of fMRI scan, to avoid the potential short-term effects of isoflurane that has been found to be most prominent during the 5 min after isoflurane cessation (Dvořáková et al., 2022).

#### AdEx mean-field model

The mean-field equations for the AdEx network are given to a first-order by Di Volo et al., 2019:

$$T\frac{d\nu_{e,i}}{dt} = F_{e,i}(W, \bar{\nu}_e, \nu_i) - \nu_{e,i}$$
$$\frac{dW}{dt} = -\frac{W}{\tau_w} + b\nu_e + a(\mu_V(\bar{\nu}_e, \nu_i, W) - E_L)$$

where $\nu_{e,i}$ is the mean neuronal firing rate of the excitatory and inhibitory population, respectively, W is the mean value of the adaptation variable, F is the neuron transfer function (i.e., output firing rate of a neuron when receiving excitatory and inhibitory inputs with mean rates $\nu_e$ and $\nu_i$ and with a level of adaptation W), a and b are the sub-threshold and spike-triggered adaptation constants, $t_w$ is the characteristic time of the adaptation variable, T is a characteristic time for neuronal response, $\mu_V$ is the average membrane voltage and $E_L$ is the leakage reversal potential. The simulations presented in this paper corresponds to the parameters: a=0; b=0 for AI state and b=300pA for SWD dynamics (b=0 for inhibitory neurons); $t_w$=200ms; T=5ms; $E_L$=-63mV for excitatory neurons and -65mV for inhibitory neurons. The main assumption of the model concerns the size of the neuronal population and the characteristic time of the neuronal dynamics. The size of the neuronal populations must be large enough to ensure the validity of a statistical description (starting in the order of thousands of neurons), and the characteristic time of the population dynamics must be slow enough to be captured by the mean-field formalism (with a lower bound in the order of the few milliseconds). These two conditions are satisfied for the system studied in this paper (for further details on the model, see Di Volo et al., 2019.

This model has been widely tested and used for the simulation of different brain states (i.e. asynchronous-irregular vs slow-waves sleep, (Goldman et al., 2023), neuronal responsiveness (Goldman et al., 2023; Di Volo et al., 2019), and whole-brain dynamics (Goldman et al., 2023). In addition, modeling tools to calculate brain signals (such as LFP and BOLD-fMRI) from this type of mean-fields have been developed (Tesler et al., 2022, 2023). For further validation, we show in Figure 6–figure supplement 2 the comparison of the SWD type of dynamics in the mean-field and in the corresponding spiking-neural network of AdEx neurons. We see that, although the amplitude of the oscillations is larger in the spiking-network, the mean-field can correctly capture the general dynamics and frequency of the SWD pattern.

#### Single region activity during stimulation in whole brain simulation

For a better illustration of the response to external stimulation during the different states we show in Figure 6–figure supplement 1, the response of single brain regions during the stimulation protocol for each state. In particular we show the stimulated area (primary visual cortex, V1) and a neighboring region (mediolateral visual area) with strong structural connectivity to V1 (see caption of the figure for details). The stimulation of a specific region is simulated as an increase in the excitatory input to the specific node. In particular, we use a square function for representing the stimulus (see panel A in Figure 6–figure supplement 1). As we can see in the figure, during the inter-ictal period (panels A, B, C) the activity is strongly driven by external stimulation and the firing rates follow the stimulation pattern. On the other hand, during the ictal period (panels D, E, F) the activity is mainly driven by the global oscillatory dynamics even during stimulation. We see in panels E, F that the stimulation can lead to a certain variation on the amplitude of the oscillations during some of the cycles, but the neuronal activity remains predominantly dominated by the global ongoing activity (SWD dynamics). This difference in responsiveness between the two states is captured in the statistical maps shown in the main text. To build the statistical maps, an ANOVA (analysis of variance) test was used. This test is originally thought to assess the significance of the change in the mean between two samples and is calculated via an F-test as the ratio of the variance between and within samples. In our case, it allowed us to assess the impact of the stimulation on the ongoing neuronal activity by performing a comparison of the timeseries of the firing rate with and without stimulation (this was performed independently for each state). For the results presented in this paper, the ANOVA analysis was performed using the “f_oneway” function of the scipy.stats. module in python.
